# Cytokine-Induced Cell Exhaustion to Mitigate Hyperinflammation

**DOI:** 10.1101/2024.08.12.607544

**Authors:** Tommaso Marchetti, Samantha Milanesi, Diana Tintor, Tiziana Lorenzini, Severin Walser, Stefano Vavassori, Onur Boyman, Jana Pachlopnik Schmid

## Abstract

Haemophagocytic lymphohistiocytosis (HLH), a life-threatening hyperinflammatory disorder often driven by dysfunctional cytotoxic CD8^+^ T cells, is marked by cytokine storms, which may follow viral infections. In a study of a perforin-deficient mouse model of HLH with a viral trigger, we aimed to determine if CD8^+^ T cell behaviour could be modulated by targeted interleukin (IL)-2 treatment. We observed a paradoxical benefit that contrasted with IL-2’s typical role in boosting T cell activity: targeted IL-2 delivery to CD8^+^ T cells led to reduced hyperinflammation and disease severity. Our results demonstrated that IL-2 induced exhaustion in overactive CD8^+^ T cells, thus mitigating hyperinflammation. These findings highlight the context-dependency of cytokine treatment and suggest new therapeutic strategies for HLH and other inflammatory diseases by leveraging cell exhaustion.

## Main

Haemophagocytic lymphohistiocytosis (HLH) is a life-threatening disorder characterized by multi-organ failure and hyperinflammation driven by overwhelming cytokine production^1^. Public awareness of HLH has been increased recently by detailed reports of SARS-CoV-2-induced hyperinflammation that demonstrated the potentially lethal nature of exaggerated immune responses^2^. While most individuals recover from viral infections through a self-limiting reaction, some patients experience a severe course of disease, a “cytokine storm” and subsequent tissue damage, with an excessive immune response to the viral infection, rather than the virus itself. In HLH, the hallmark of pathology is defective cytotoxicity of the CD8^+^ T cells that usually eliminate virus-infected cells via perforin (Prf)-mediated killing^3^. This defective cytotoxicity results in an unbalanced immune response^4–7^ in which the exaggerated release of proinflammatory cytokines^8–10^ is due to ineffective killing and thus prolonged interactions between CD8^+^ T cells and their targets ^11^. These processes ultimately lead to HLH.

The cytokine interleukin (IL)-2 is essential for T cell survival, proliferation and function^12^. IL-2 has long been recognized for its critical role in activating the effector arm of the immune system, particularly in enhancing the activity of cytotoxic T cells ^13^. However, subsequent research revealed that IL-2’s effects are highly context-dependent and promote both inflammatory and anti-inflammatory outcomes^14–17^. The development of antibodies that selectively target IL-2 to specific T cell subpopulations has greatly advanced our understanding of the cytokine’s diverse actions^18–20^. These antibodies can direct IL-2 to its trimeric receptor, biased towards the receptor’s α-chain (CD25) or its dimeric receptor, biased towards the receptor’s β-chain (CD122). This targeted approach shows great promise for treating autoimmune diseases by directing IL-2 to the trimeric receptors present on virtually all regulatory T cells (Tregs)^21,22^ and for treating cancer by directing IL-2 to cells that predominantly express the dimeric IL-2 receptor, such as CD8^+^ T cells^23,24^. However, the application of these strategies and the role of IL-2 in hyperinflammatory conditions like HLH have not been extensively characterized. Attempts in mouse models to directly enhance Treg activity via IL-2 manipulation have not only failed to relieve HLH but have even exacerbated the disease^25^.

The present study uncovered a surprising and paradoxical benefit of targeting CD8^+^ T cells with IL-2. These findings may provide new insights into how promoting CD8^+^ T cell activity and subsequent exhaustion could be advantageous in treating HLH, a disease traditionally associated with the harmful effects of these cells.

## Results

### *In vivo* delivery of a CD122-biased IL-2 complex counteracts hyper-inflammation and improves health in mice with an impairment in the cytotoxic machinery

To examine how the delivery of IL-2 to different receptors affects HLH severity, we employed a targeted approach using IL-2/anti-IL-2 antibody complexes in a Prf-deficient (Prf1KO) mouse model of HLH that effectively replicates the signs of the human disease^9^.

Following infection with the WE strain of lymphocytic choriomeningitis virus (LCMV), mice were treated for three consecutive days with either the IL-2/JES6-1 complex, referred to as JES6-1 and which preferentially directs IL-2 to the trimeric IL-2 receptor^18,26^, or the IL-2/NARA1 complex, referred to as NARA1, which targets IL-2 to cells expressing the dimeric IL-2 receptor^24^ (Fig. 1a). We analysed the mice on post-infection day 9 or post-infection day 15, i.e. one day or one week after treatment (Fig. 1b). Administration of JES6-1 resulted in 100% mortality (Fig. 1c). In contrast, NARA1 treatment did not reduce the survival rate and significantly improved the overall health of the Prf1KO mice, as evidenced by a temporary weight gain (Fig. 1d) and an alleviation of key signs of HLH, such as splenomegaly and anaemia (Fig. 1e). The total blood cell counts remained within the ranges observed in wild-type (WT) mice (Fig. 1f). Splenomegaly was also less pronounced in WT mice, suggesting immune system benefits beyond those seen in Prf1KO mice (Fig. 1e). The reduced spleen size and cellularity even persisted one week after treatment (Extended Data Fig. 1b). NARA1 therapy altered the serum cytokine profile, with lower levels of IFNγ, a key driver of HLH^9,10^, and higher levels of the anti-inflammatory cytokine IL-10^27,28^ and TNFα in infected Prf1KO mice (Fig. 1g). The treatment did not affect ferritin levels (Fig. 1h and Extended Data Fig. 1f) but was associated with greater expression of IL-2 receptor a (Extended Data Fig. 1d).

**Fig. 1.**
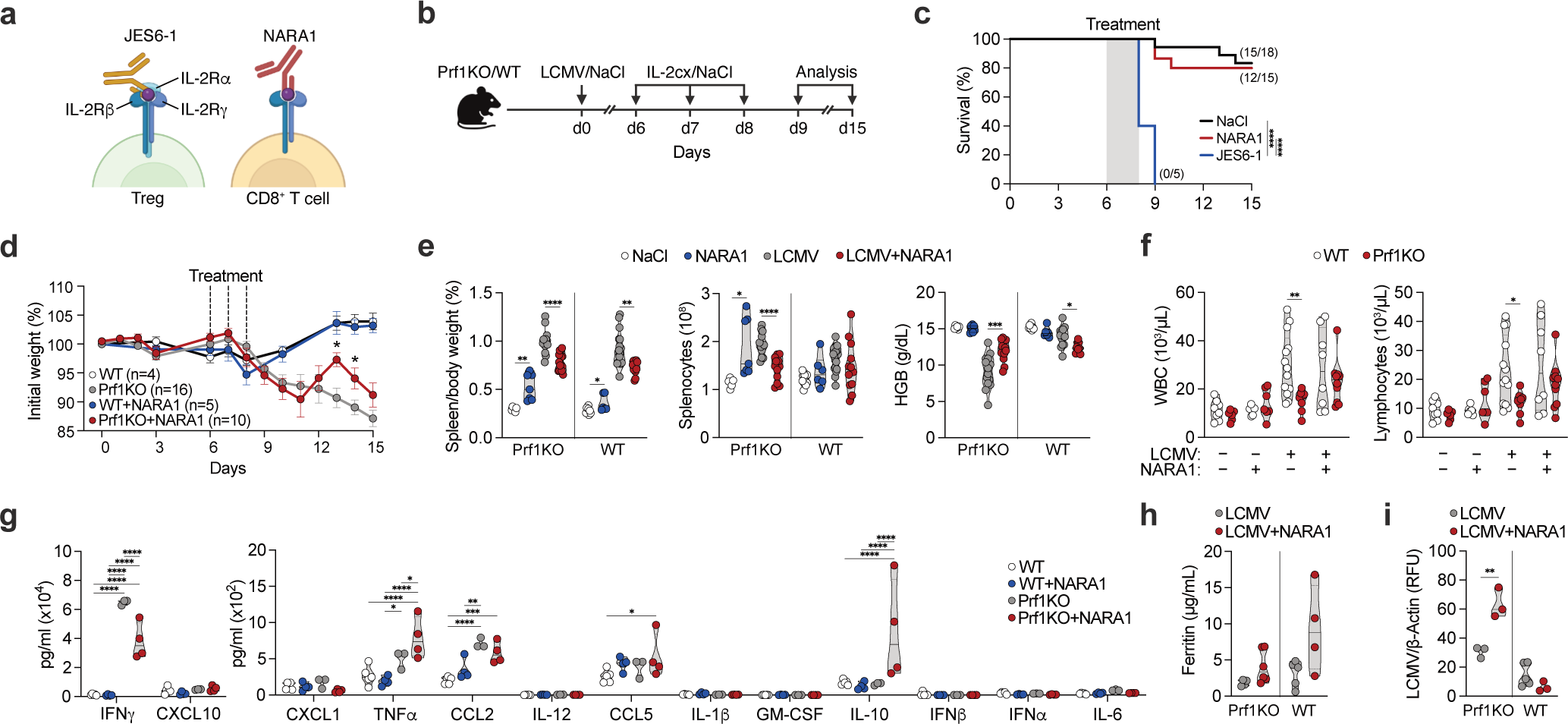
*In vivo* delivery of CD122-biased IL-2 complex counteracts hyperinflammation and improves health in mice with an impairment in cytotoxicity. **a,** Illustration of IL-2 complexes binding to receptors. **b**, The experimental set-up: WT and Prf1KO mice received an injection of saline solution or 200 pfu of LCMV-WE on day 0, followed by injections of saline solution or IL-2 complexes for three consecutive days (post-infection days 5 to 7). Spleen and blood analyses were conducted on post-infection days 9 and 15. **c**, Kaplan-Meier survival curves of Prf1KO mice after infection with LCMV-WE: the numbers in brackets correspond to the survivors at the endpoint and the total starting population. **d**, The changes in the mean _±_ standard error body weight after infection with LCMV-WE. A mixed-effect analysis with Šidák’s multiple comparison test was applied. **e**, The variables of the spleen-to-mouse weight ratio, the splenocyte count, and the haemoglobin (HGB) level on post-infection day 9. **f,** White blood cell (WBC) and lymphocyte counts in mice on post-infection day 9. **g,** Serum levels of cytokines involved in the mouse antiviral response on post-infection day 9. A two-way analysis of variance with Tukey’s correction was performed. **h,** The serum ferritin concentration on post-infection day 9. **i**, Splenic viral titres on post-infection day 9. RFU=relative fluorescent unit. Each dot represents a biological replicate pooled from two or more independent experiments. n=5 to 18 per group (**e**); n=6 to 13 per group (**f**); n=3 to 5 per group (**g** and **i**); n=4 to 5 per group (**h**). Truncated violin plots with the quartiles, range, and mean are shown alongside scatter plots (**e−h**). If not specified otherwise, two-tailed t-tests were applied; the statistical significance of the result is indicated by **** p<0.0001, *** p<0.001, ** p<0.01, and * p<0.05.

We initially hypothesized that directing IL-2 to CD8^+^ T cells would enhance their perforin-independent killing mechanisms and control viral infection. However, NARA1 treatment of Prf1KO mice unexpectedly increased the viral load in spleens on day 1 post-treatment (Fig. 1i). The observed changes in blood cytokine levels, cell counts, and the negative impact on viral control were temporary and disappeared one week after the cessation of NARA1 treatment (Extended Data Fig. 1c,e and g). These findings indicate that the transient anti-inflammatory effect and health improvement seen in NARA1-treated Prf1KO mice resulted from mechanisms unrelated to viral clearance.

### During an infection, the CD122-biased IL-2 complex selectively modulates CD8^+^ T cell differentiation independently of Prf

To explore the mechanism by which NARA1 exerted its anti-inflammatory effect in our murine model of HLH, we investigated cell populations potentially influenced by the administration of CD122-biased IL-2 complexes. While these complexes are known to primarily expand CD8^+^ T cells, emerging evidence suggests that the complexes also have an impact on other immune cell populations, such as Tregs, natural killer (NK) cells and dendritic cells (DCs)^29^.

While NARA1 was associated with a relative increase in the number of Tregs in uninfected mice on post-treatment day 1, this effect was less pronounced in the LCMV infection model (Fig. 2a). Additionally, NARA1 did not affect the proliferation capacity of Tregs or the total splenic Treg count during the infection (Fig. 2b and Extended Data Fig. 2a). Previous studies have indicated that NK cells can regulate CD8^+^ T cell activity by impacting viral clearance and the inflammatory state^30–32^. In non-infected mice, NARA1 treatment was associated with significantly higher relative and absolute NK cell counts in the spleen, as well as increased NK cell proliferation (Fig. 2c, d and Extended Data Fig. 2b). However, after LCMV infection, the splenic NK cell counts were low and decreased further following NARA1 administration (Fig. 2c). Although IL-2 treatment can indirectly increase the number of dendritic cells, which are crucial for antiviral immunity^33,29^, we did not observe this effect under any of our experimental conditions (Extended Data Fig. 2c). These findings indicate that although NARA1 affects the counts and proliferation of Tregs, NK cells and DCs under certain conditions, these changes do not fully account for the observed anti-inflammatory effects.

**Fig. 2.**
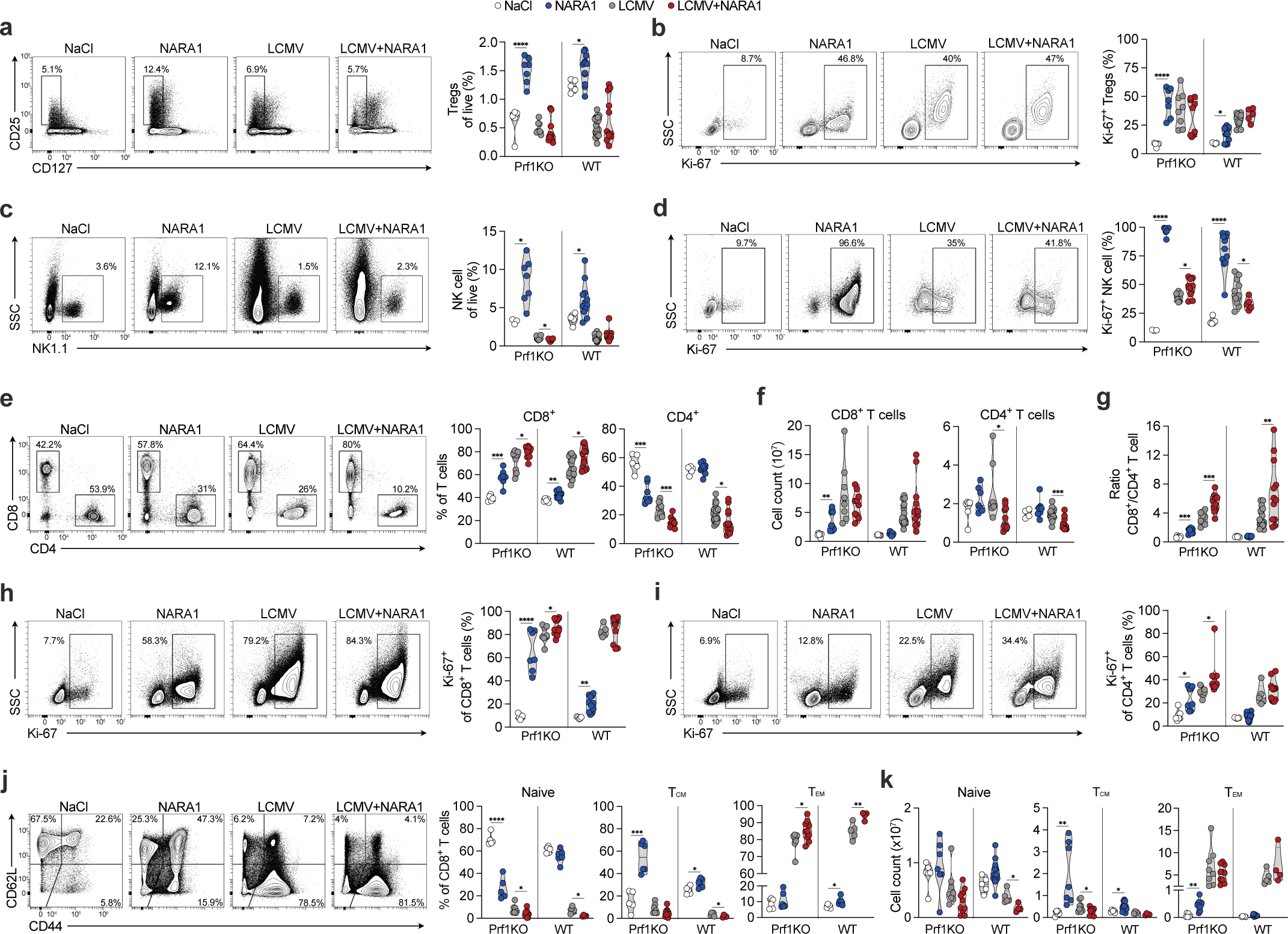
CD122-biased IL-2 selectively modulates CD8 T Cell differentiation independently of Prf, during infection. **a**, Representative flow cytometry plots (left) and counts (right) of Tregs gated on CD4^+^ T cells. **b**, Flow cytometry plots (left) and counts (right) of Ki-67 expression by Tregs. **c**, Flow cytometry plots (left) and counts (right) of splenic NK cells gated on lymphocytes. **d**, Flow cytometry plots (left) and counts (right) of Ki-67 expression by NK cells. **e**, Flow cytometry plots (left) and counts (right) of relative CD8^+^ and CD4^+^ T cells. **f**, Total count of CD8^+^ and CD4^+^ T cells. **g**, Ratios of CD8^+^ T cells and CD4^+^ T cells. **h**, Flow cytometry plots (left) and counts (right) of Ki-67 expression by CD8^+^ T cells. **i**, Flow cytometry plots (left) and counts (right) of Ki-67 expression by CD4^+^ T cells. **j**, Flow cytometry plots (left) and counts (right) illustrating relative levels of CD8^+^ T cell subsets: naive (CD62L^+^, CD44^−^), central memory (T_CM_) (CD62L^+^, CD44^+^), and effector memory (T_EM_) (CD62L^−^, CD44^+^). **k**, CD8^+^ T cell subset counts. All the cells were taken from spleens on post-infection day 9. All the representative flow cytometry plots come from Prf1KO mice. Each dot on the scatter graph represents a biological replicate pooled from two or more independent experiments. n=5 to 13 per group (**a**), 5 to 12 per group (**b**,**h**,**i**,**k**), 3 to 18 per group (**c**), 3 to 12 per group (**d**), 5 to 18 per group (**e**−**g**), and 5 to 10 per group (**j**). Truncated violin plots with the quartiles, range, and mean are shown alongside scatter plots. Two-tailed t-tests were applied; the statistical significance of the result is indicated by **** p<0.0001, *** p<0.001, ** p<0.01, and * p<0.05 (p>0.05 in the absence of asterisks).

Consequently, we next investigated the impact of NARA1 on conventional T-cell populations. NARA1 treatment resulted in a relative increase in CD8^+^ T cells, regardless of the Prf deficiency or infection status (Extended Data Fig. 2e,d). However, this relative increase did not translate into greater absolute splenic CD8^+^ T cell counts (Fig. 2f). Conversely, NARA1 treatment was associated with lower relative and absolute CD4^+^ T cell counts (Fig. 2f), suggesting a preferential expansion of CD8^+^ T cells (Fig. 2g). Analysis of proliferating cells using Ki-67 staining indicated enhanced proliferation within the CD8^+^ T cell compartment compared to the CD4^+^ T cell compartment (Fig. 2h,i). The impact of NARA1 on CD8^+^ T cell maturation varied with the timing of the administration, relative to the start of the infection. In uninfected mice, NARA1 treatment mainly induced CD8^+^ T cells to take on a central memory (T_CM_) (CD44^+^, CD62L^+^) phenotype. During LCMV infection, however, NARA1 treatment shifted the CD8^+^ T cell population towards a predominantly effector memory (T_EM_) (CD44^+^, CD62L^−^) phenotype, regardless of the cytotoxic capacity (Fig. 2j). Despite this shift, the overall number and proliferation stage of CD8^+^ T_EM_ cells remained unchanged (Fig. 2k and Extended Data 2e). A week after treatment, relative CD8^+^ T cell counts were higher in NARA1-treated mice, although no differences in the T_EM_ compartment were observed (Extended Data Fig. 2f,g).

In line with the results of previous reports on the use of CD122-biased complexes to deliver IL-2 to CD8^+^ T cells^18,24^, we found that NARA1 treatment also delivers IL-2 to these cells in the context of hyperinflammation. Our results challenge the hypothesis whereby exaggerated IL-2 signalling in CD8^+^ T cells primarily fuels HLH and, intriguingly, suggest that the signalling is even potentially advantageous in this scenario.

### Transcriptomic analysis reveals the induction of an exhausted signature in Prf-deficient CD8^+^ T cells following IL-2 stimulation

IL-2 is known to stimulate the immune system to react to viruses and tumours^34,35^ but can also induce early exhaustion^36,37^; this early exhaustion has been considered a mechanism for preventing fatal immune diseases ^38,39^. In our study, we observed that during infection, NARA1 treatment expanded and differentiated CD8^+^ T cells regardless of their cytotoxicity. However, in the Prf1KO model, NARA1 treatment led to a decrease in the overall immune response, as evidenced by lower IFNγ levels (Fig. 1g) and higher viral titres (Fig. 1h).

To better understand the impact of IL-2 on CD8+ T cells lacking cytotoxic function, we used next-generation RNA sequencing (seq) to analyse and compare cells isolated from infected NARA1-treated mice and untreated Prf1KO mice on post-infection day 9. In line with our previous results, pathways involving immune system restraint were the most upregulated in treated mice (Fig. 3a), while T cell activation pathways were mostly downregulated (Extended Data Fig. 3a). NARA1 treatment in Prf1KO mice led to the upregulation of genes linked to effector functions (*Klrb1c, Tbx21, Gzma, Gzmb, Gzmk, Fasl, Id2*) and genes associated with exhaustion (*Pdcd1, Tim-3, Lag3, Tigit, Cd244a*). In contrast, genes typically present in the earlier stages of CD8^+^ T cell development or associated with a memory-like phenotype (*Tcf7, Id3, Eomes, Ikzf2*) were downregulated (Fig. 3b). By post-infection day 15, we observed a reversal of the upregulated exhaustion signature in Prf1KO mice noted on post-infection day 9 (Fig. 3c); again, this was in line with the previously observed transient effect of NARA1.

**Fig. 3.**
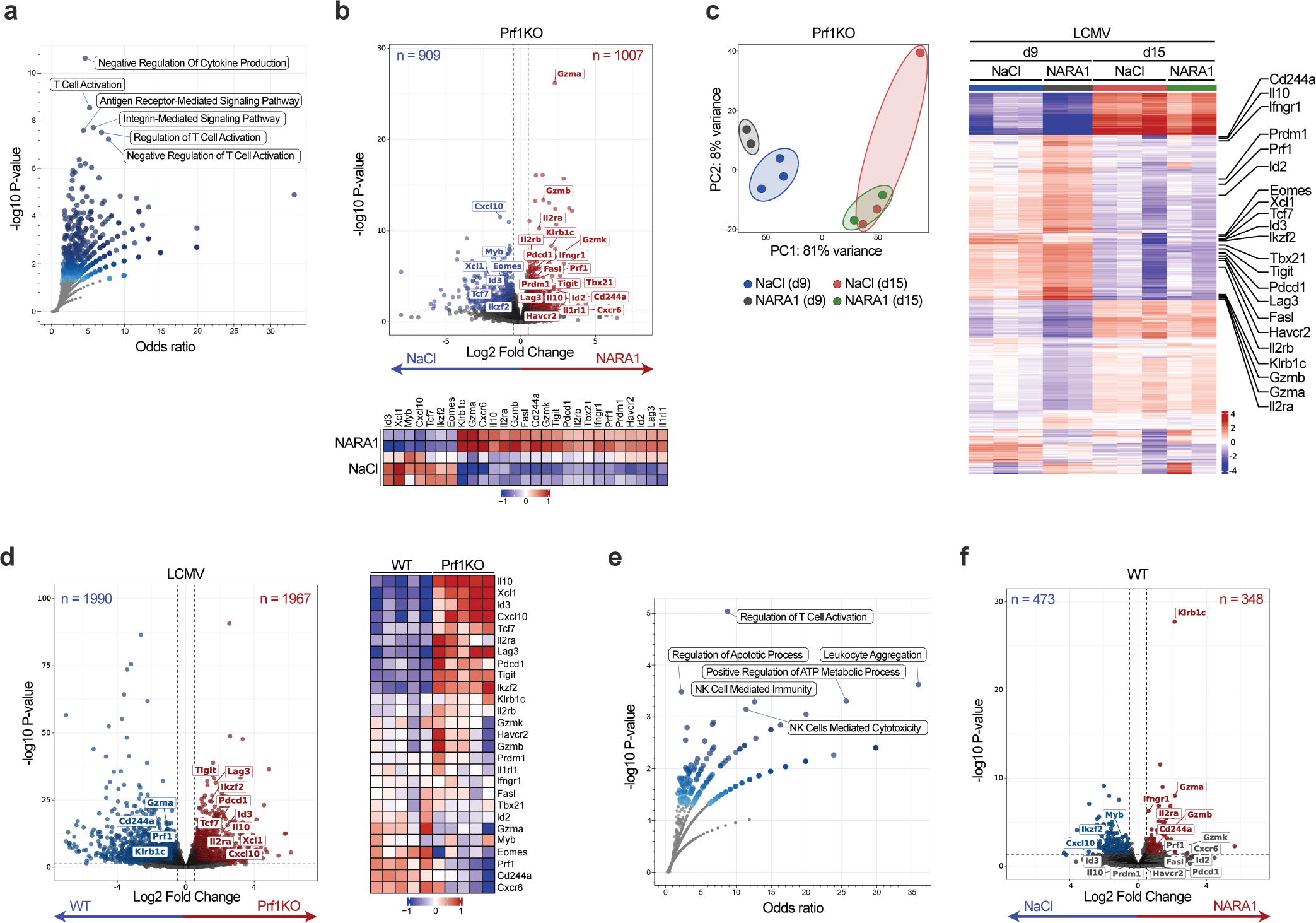
Transcriptomic analysis reveals induction of an exhausted signature in Prf-deficient CD8 T cells following IL-2 stimulation. **a,** Dot plot of the results of a Gene Ontology (GO) term enrichment analysis of differentially expressed genes (DEGs) upregulated in CD8 T cells from LCMV-infected Prf1KO mice treated with IL-2 (saline solution n=3, NARA1 n=2). **b**, CD8+ T cells obtained from the spleens of LCMV-infected mice under various conditions were analysed using RNAseq. The volcano plot (top) displays upregulated and downregulated genes in CD8^+^ T cells from Prf1KO mice treated with IL-2, compared with non-treated mice on post-infection day 9. The heatmap (bottom) illustrates the mean relative expression of specific genes. **c**, A principal component analysis (left) of RNAseq data from splenic CD8^+^ T cells generated on post-infection days 9 and 15. The heat map (right) shows the top DEGs for the various treatment conditions and time points. **d**, The volcano plot (left) displays upregulated and downregulated genes in CD8^+^ T cells from Prf1KO mice, compared with WT mice on post-infection day 9. The heatmap (right) shows the mean relative expression of the genes illustrated in (B). **e**, A dot plot of GO term enrichment analysis of DEGs upregulated in CD8^+^ T cells from LCMV-infected WT mice treated with IL-2. **f**, The volcano plot displays genes that are upregulated and downregulated in CD8^+^ T cells from LCMV-infected WT mice treated with IL-2, compared with non-treated mice on post-infection day 9 (saline solution n=5, NARA1 n=4).

RNAseq of samples from LCMV-infected WT and Prf1KO mice revealed a pre-existing, activated state in CD8^+^ T cells with impaired cytotoxic function. These T cells also displayed an exhaustion signature, as characterized by the elevated expression of the inhibitory receptors Tigit, Lag3 and Pdcd1 and the *Il2ra* gene coding for the IL-2 receptor a(Fig. 3d). We observed that *Il10* gene expression was elevated in infected Prf1KO mice and then further amplified by NARA1 treatment (Fig. 3b,d). These findings fit with the observed elevation in serum IL-10 levels (Fig. 1g) and point to a potential role of CD8^+^ T cells in the secretion of this cytokine. Our observations indicated that in the context of HLH, providing CD8^+^ T cells with IL-2 may exacerbate their pre-existing overactivated state. To investigate NARA1’s effect on Prf-proficient CD8^+^ T cells, we compared the transcriptomes of treated vs. non-treated WT mice on post-infection day 9. Gene enrichment analysis showed increased activity in T cell activation pathways (Fig. 3e and Extended Data Fig. 3b), along with genes related to an effector signature such as *Gzma* and *Gzmb* and *Il2ra* in NARA1-treated mice (Fig. 3f). In contrast to the signature seen in Prf1KO mice, genes tied to exhaustion were not upregulated.

These findings indicate that administering IL-2 to overactivated, Prf-deficient CD8^+^ T cells might initiate a “shutdown” program that potentially accounts for the observed reduction in the signs of immunopathology.

### IL-2 prompts CD8^+^ T cells to adopt an exhausted phenotype and functional state essential for mitigating hyper-inflammation

T-cell exhaustion has long been considered to be a harmful event that must be reversed for restoration of the immune system’s ability to fight infections and kill tumour cells^40–42^. However, this exhaustion mechanism probably evolved to prevent the immune system from becoming hyperactive and damaging self-tissues^43,44^. This possibility suggests that the T cells’ intrinsic exhaustion program could be exploited to dampen immunopathology.

To determine how NARA1 treatment sensitizes CD8^+^ T cells to exhaustion, we initially used high-parameter spectral flow cytometry to simultaneously evaluate surface receptors and transcription factors on splenocytes. The results were visualized with uniform manifold approximation and projection (UMAP). NARA1 administration altered the distribution and marker expression of CD8^+^ T cells in LCMV-infected Prf1KO mice (Fig. 4a). A PhenoGraph analysis^45^ of marker expression patterns (Fig. 4b and Extended Data Fig. 5a) revealed eight distinct CD8^+^ T cell clusters (Fig. 4c): “naïve” CD8^+^ T cells characterized by TCF-1 and CD62L expression; T_CM_ characterized by TCF-1 and CD44 expression; “precursor exhausted T cells” (T_PEX_) characterized by TCF-1 and PD-1 expression; “T follicular helper” CD8^+^ T cells (T_FH_) characterized by BCL-6 expression; “short-lived” T_EM_ characterized by CD44 and KLRG1 expression, and three T_EM_ clusters characterized by CD44, CD183, CD122, and PD-1 expression. A subsequent evaluation with the “significance analysis of microarrays” method^46^ indicated that NARA1 expanded the T_EM_ population, as characterized by high levels of CD122 and PD-1 expression (Fig. 4d). Notably, NARA1 treatment appeared to constrain the T_PEX_ and short-lived T_EM_ cell populations. These effects were not observed in WT mice (Extended Data Fig. 4a−d). However, the reduction in the naïve cluster upon NARA1 administration indicated that providing cytotoxicity-proficient CD8^+^ T cells with IL-2 induced differentiation without increasing the expression of genes characteristic of exhaustion (Fig. 3f).

**Fig. 4.**
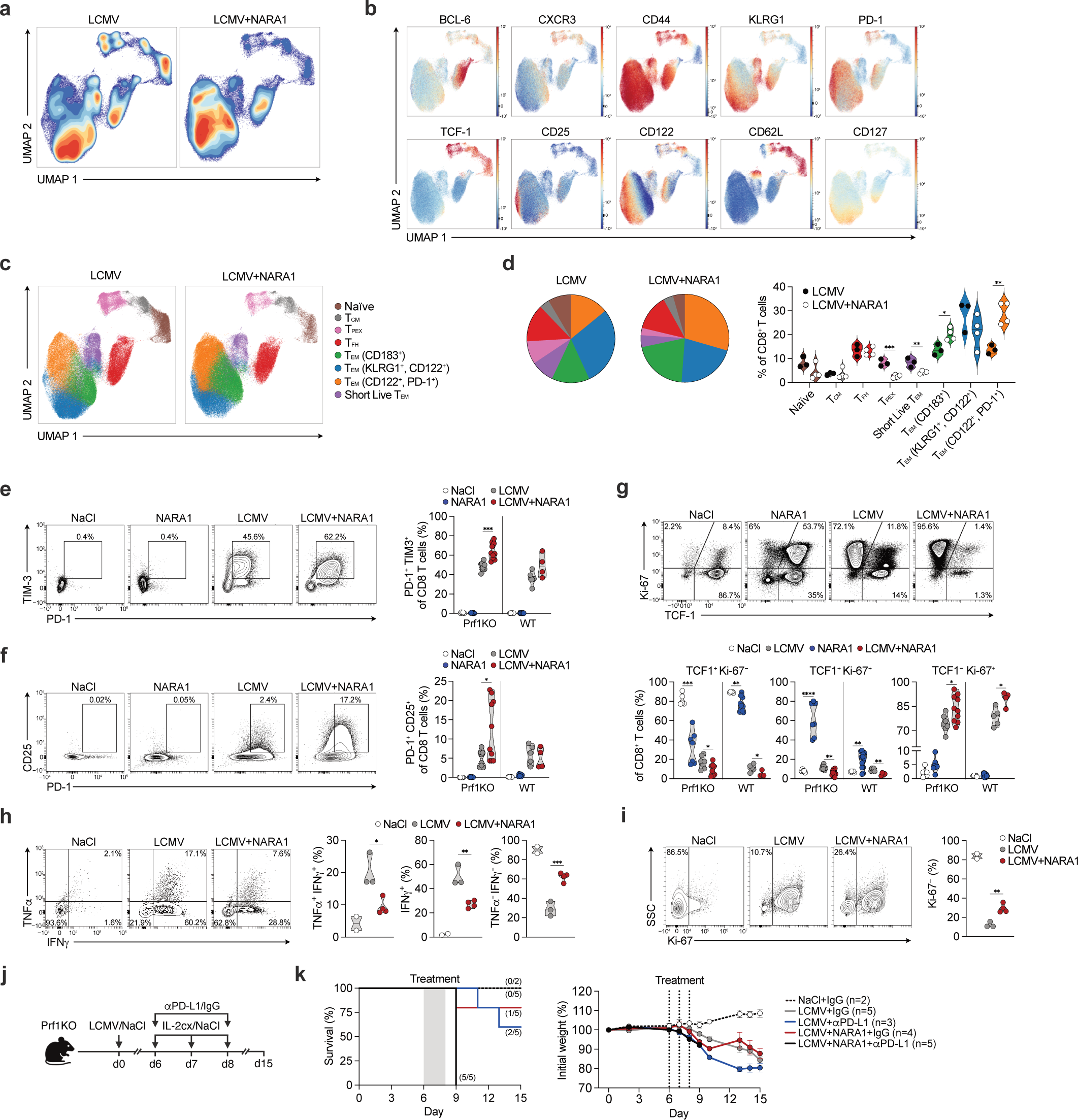
*In vivo* IL-2 treatment prompts Prf1KO CD8 T cells to adopt an exhausted phenotype and a functional state that is essential for mitigating hyperinflammation. **a,** Stacked uniform manifold approximation and projection (UMAP) plots of high-dimensional spectral flow cytometry data from gated CD8^+^ T cells obtained from the spleen of LCMV-infected Prf1KO mice after treatment with LCMV (n=3) or LCMV+NARA1 (n=4) on post-infection day 9, coloured by distribution density. **b**, UMAP plots of selected markers in cells from Prf1KO mice infected with LCMV and treated with NARA1, scaled by fluorescent intensity. **c**, UMAP plots of CD8^+^ T cells from Prf1KO infected mice, subclustered using PhenoGraph and 26 marker expression profiles. T_CM_=central memory T cells, T_PEX_=pre-exhausted T cells, T_FH_=T follicular helper cells, T_EM_=effector memory T cells. **d**, Proportions of cells, by treatment. **e**, Flow cytometry plots (left) and counts (right) of PD-1 and TIM3 double-positive CD8^+^ T cells. **f**, Flow cytometry plots (left) and counts (right) of PD-1 and CD25 double-positive CD8^+^ T cells. **g,** Flow cytometry plots illustrating the relative counts of CD8^+^ T cell subsets expressing TCF-1 and Ki-67. **h**, Flow cytometry plots and counts depicting IFNγ and TNFα production by CD8^+^ T cells from spleens collected on post-infection day 9, following stimulation for 5 h. The plots were gated on CD8^+^ T cells. **i**, Flow cytometry plots and counts depicting Ki-67 expression by splenic CD8^+^ T cells, following stimulation for 5 h. The plots were gated on CD8^+^ T cells. **j**, Experimental set-up: Prf1KO mice received injections of 200 pfu of LCMV-WE or saline solution, followed by subsequent injections of IL-2 complexes or saline solution and/or 200 µg of anti-PD-L1 or IgG. **k**, Survival curves (left) and mouse weight (right) following infection. The numbers in brackets correspond to the survivors at the endpoint and the total starting population. Each dot represents a biological replicate pooled from two or more independent experiments, except for (**h**) and (**i**). n= 5 to 12 per group (**e** and **f**), 5 to 13 per group (**g**), and 2 to 4 per group for (**h** and **i**). Truncated violin plots with the quartiles, range, and mean are shown alongside scatter plots. Two-tailed t-tests were applied; the statistical significance of the result is indicated by **** p<0.0001, *** p<0.001, ** p<0.01, and * p<0.05.

In line with RNAseq results, flow cytometry experiments revealed a greater prevalence of CD8^+^ T cells that were double-positive for the PD-1 and TIM-3, markers of severe T cell exhaustion^47^, in NARA1-treated Prf1KO mice during an infection (Fig. 4e). Previous literature has shown that IL-2 stimulation promotes CD25 upregulation on T cells^36^. Consistently, we observed the simultaneous upregulation of CD25 and PD-1, along with the downregulation of TCF-1, in Prf1KO mice (Fig. 4f and Extended Data Fig. 4f). Following an infection, CD8^+^ T cells typically expand while their ability to self-renew, as indicated by the TCF-1 marker, diminishes^48^. NARA1 treatment of Prf1KO mice appeared to accelerate the loss of TCF-1, as also shown by decreased expression of *Tcf7* (Fig. 3b). Furthermore, we observed greater proliferation of TCF-1^−^ CD8^+^ T cells, which suggests that NARA1 treatment may not preserve the stem-like cell population (Fig. 4g). Our data also indicate that NARA1 treatment decreases the population of long-lived effector (KLRG1^+^, CD27^−^) T cells^49^ (Extended Data Fig. 4e); although *Fasl* mRNA expression was upregulated (Fig. 3b), a corresponding increase in FASL or TRAIL expression was not detected (Extended Data Fig. 4g).

To assess the functional impact of NARA1-induced exhaustion on *Prf1*-deficient CD8^+^ T cells, we analyzed their cytokine production and proliferation after restimulation. NARA1 treatment was associated with a significantly reduced ability of CD8^+^ T cells to produce effector cytokines, such as IFNγ and TNFα (Fig. 4h), as well as an impairment in their proliferative capacity (Fig. 4i). A week after treatment, however, we found that both functions had been restored (Extended Data Fig. 4h,i). To test the hypothesis that inducing exhaustion in effector cells may be beneficial in HLH models and to examine the exhaustive effect of NARA1 on cytotoxicity-deficient CD8^+^ T cells during infection, we used immune checkpoint blockade to reverse exhaustion (Fig. 4j). Mice receiving both NARA1 and anti-PD-L1 treatment succumbed within a day of treatment, whereas most of the mice receiving either anti-PD-L1 alone or NARA1 alone survived (Fig. 4k).

Our results suggest that IL-2 delivery to CD8^+^ T cells harnesses a controlled form of exhaustion required to mitigate the overactive immune response in HLH, potentially reduce tissue damage, and maintain a level of immune surveillance that may be critical for managing the immune disease.

## Discussion

IL-2 exhibits strongly contrasting dual effects on the immune system. While IL-2 acts as a potent immunostimulator for CD8^+^ T cells by promoting their activation and proliferation and thus enhancing their ability to combat infections and tumours^13,36^, it can also suppress immune responses by fostering the activity of Tregs^15^. However, this potential Treg-mediated dampening effect might help to mitigate the development of autoimmune diseases and graft-versus-host disease^50^.

In the present study, we explored cytokine manipulation as a possible therapeutic strategy for HLH. Our results showed that the delivery of IL-2 by cytokine-antibody complexes to dysfunctional CD8^+^ T cells may induce exhaustion, a natural process that limits T cell immune responses and thereby provides therapeutic benefits in mouse models of the disease. Our findings are in line with previous reports that IL-2 can induce exhaustion and reduce the CD8^+^ T cell count when administered during the cell expansion phase^36,37^. Notably, treatment with NARA1, a CD122-biased IL-2 delivery system, alleviated several core signs of HLH, including splenomegaly, cytopenia, and low haemoglobin levels, and resulted in a transient decrease in levels of IFNγ, the key driver of HLH. We found that NARA1-mediated IL-2 delivery suppressed TCF-1 expression and promoted an effector differentiation state in CD8^+^ T cells. This progression drives CD8^+^ T cells towards a hyper-activated and eventually exhausted phenotype and thus reduces their ability to produce inflammatory cytokines. We believe that this mechanism underlies the observed clinical improvements. CD8^+^ T cells expressing inhibitory receptors are known to secrete the suppressive cytokine IL-10^47^. Following NARA1 administration, we observed elevated IL-10 levels in the blood and in CD8^+^ T cell RNA. This suggests that IL-10 production by CD8^+^ T cells may function as a self-regulatory mechanism to mitigate excessive cell-mediated immunopathology during hyperinflammation, as previously proposed^51^. Furthermore, it would be interesting to investigate whether IL-2 initiates this regulatory process.

Intriguingly, recent investigations of this topic have given contrasting results. Administration of IL-2/NARA1 complexes and an IL-2–NARA1 complex fusion protein under non-infection conditions was associated with lower levels of exhaustion markers^24,52^. This discrepancy prompts speculation about IL-2’s differential effects on CD8^+^ T cells, depending on their activation state (effector vs memory phases). It is plausible that these observed differences arise from the distinct immunological contexts in which IL-2 is administered^53^. These differences highlight the complexity of T cell regulation and the need for further exploration of the precise mechanisms governing IL-2-mediated effects on T cell exhaustion and immune function.

The exact mechanism by which CD25- and CD122-biased IL-2 complexes yield contrasting effects in HLH remains unclear. While the targeting of differentiated CD8^+^ T cells by CD122-biased complexes appears to be therapeutically beneficial in the mouse model studied here, the administration of CD25-biased complexes, typically associated with Treg expansion, exacerbated the disease. CD25 is expressed on the surface of terminal-effector CD8^+^ T cells^54^, and reportedly reverses T cell exhaustion^55^. One can hypothesize that in the hyper-inflammatory setting of HLH, IL-2 signalling through the trimeric receptor in activated CD8^+^ T cells leads to unintended, pro-inflammatory effects. The reason why the same cytokine exhibits such contrasting functions remains elusive, but we could speculate that the immune system has evolved to respond to different ranges of IL-2 concentrations. At low concentrations, IL-2 supports Tregs, which have high-affinity IL-2 receptors, allowing them to mitigate overreactions to minor threats. As IL-2 production increases, the focus shifts to activating CD8^+^ T cells, which have lower-affinity IL-2 receptors, to combat more severe infections. However, when supraphysiological or exogenous IL-2 is administered to already overactivated T cells, as observed in our HLH model, it may induce cell exhaustion as a means of preventing severe immunopathology.

Despite these uncertainties, the targeted manipulation of immune cells through cytokines and cytokine-antibody complexes may be able to elicit effective anti-inflammatory responses. Our present results offer a new view of how dysfunctional immune cell responses might be regulated and suggest that the induction of exhaustion could be a valuable therapeutic strategy in diseases characterized by hyperactive immune reactions.

## Methods

### Mice

C57BL/6J and C57BL/6-Prf1tm1Sdz/J mice were acquired from Jackson Laboratories. Male mice aged 8 to 12 weeks were utilized in the study. Mice were randomly assigned to experimental groups for all experiments, including control and treatment conditions. Blinding was applied during weight measurement, splenocyte and serum analysis, and blood count assessments. Operators needed to be aware of the experimental conditions for other procedures to ensure proper execution. Sample sizes are specified in the figure legends, and no statistical methods were used to predetermine these sizes. The animals were housed in specific-pathogen-free facilities at the University of Zurich, with a light cycle from 6:00 to 18:00, a dark cycle from 18:00 to 6:00, room temperatures between 20 and 22 °C, and humidity levels between 30% and 70%. All experiments were approved and conducted in compliance with the guidelines and regulations of the Laboratory Animal Service Center at the University of Zurich and the Canton of Zurich Ethical Committee.

### Viral infection and treatments

Mice were infected intraperitoneally (i.p) with 200 plaque-forming unit of LCMV-WE as already described ^4^, the virus was obtained from P. Aichele (Medical Center-University of Freiburg, Freiburg, Germany). For experiments involving the IL-2/NARA1 complex (referred to as NARA1), 1.5 µg human IL-2 (Proleukin, Roche) was mixed with 15 µg NARA1 antibody in PBS, for 10 minutes at room temperature, as previously described ^24^. For experiments involving the IL-2/JES6-1 complex (referred to as JES6-1), 1.5 µg mouse IL-2 (PrepoTech) was mixed with 15 µg NARA1 antibody in PBS, for 10 minutes at room temperature, as described elsewhere ^18,26^. Complexes were injected i.p. in a total volume of 100 µl per injection to animals according to the study design. In immune checkpoint blockade experiments, mice received i.p. injections of either 200 µg of anti-PD-L1 (BioXcell) antibody or 200 µg hamster IgG isotype control (BioXcell). The timing and frequency of injections are indicated.

### Tissue processing

Mice were euthanized with CO2, after which spleens were collected and weighed before preparing the cell suspension. Single-cell suspensions of splenocytes were created by mechanically disaggregating the spleens through a 70 µm cell strainer (Corning), followed by red blood cell lysis with a lysis buffer (Merck). Blood was drawn from the heart into heparin tubes and stored at 4°C.

### Blood count and cytokine quantification

200 µl of blood was sent to the Laboratory of Animal Veterinary (Universitäres Tierspital Zürich, Zurich, Switzerland) for hematological analysis. The remaining blood was centrifuged for 10 minutes at 1200 rcf, the supernatant serum was collected and stored at −80 °C for future analysis. Serum cytokine quantification was performed using a LEGENDplex (BioLegend) assay and serum concentration was diluted at 1:2 ratio as described in the manufacturer protocol. Data acquisition was performed using a Aurora (Cytek), and analysis was performed using QOGNIT software (BioLegend). ELISA kits were used to quantified ferritin (Christal Chem) and soluble mouse CD25/IL-2 alpha receptor (R&D Systems) following manufactorers instructuon.

### Ferritin and IL-2 alpha receptor quantification

ELISA kits were used to quantified serum ferritin (Christal Chem) and soluble mouse CD25/IL-2 alpha receptor (R&D Systems) following manufacturers instruction.

### Virus titer

RNA was extracted from spleen tissue preserved in Trizol (Invitrogen) using the RNA isolation mini kit (Invitrogen), followed by conversion into cDNA using SuperScript Vilo Master Mix (ThermoFisher). Quantitative PCR was conducted on the cDNA with Fast Start SYBR Green Master mix (Roche) and these primers: LCMV forward 5′-TCTCATCCCAACCATTTGCA-3′, LCMV reverse 5′-GGGAAATTTGACAGCACAACAA-3′; β-actinforward 5′-CCAGCAGATGTGGATCAGCA-3′, and β-actin reverse 5′-CTTGCGGTGCACGATGG-3′. The expression of LCMV DNA was normalized against β-actin levels. Fluorescence measurement during PCR was performed on a CFX384 Touch Real-Time PCR System (Bio-Rad) and analysed using CFX Maestro software (Bio-Rad).

### Cell phenotyping and stimulation

Cells were stained with specified surface antibodies and ZombieAqua (BioLegend) for 30 minutes at 4°C in PBS containing 4% FBS and 0.01% NaN_3_. For intracellular staining, cells were processed using a FoxP3 transcription factor staining kit (eBioscience) and then incubated overnight at 4 °C with intracellular antibodies. For detecting intracellular cytokines, cells were suspended in RPMI supplemented with 5% heat-inactivated fetal bovine serum, 2 mM L-glutamine, 10 mM Non-Essential Amino Acids Solution and 100 U/ml penicillin-streptomycin. The cells were then placed into the wells of a 96-well plate, and medium containing a cell stimulation cocktail plus protein transport inhibitors (eBioscience) was added following manufacturer instruction, for a 5-hour incubation at 37 °C. Following this incubation, cells underwent extracellular staining, were fixed and permeabilized, and subsequently stained for intracellular markers. Data collection was carried out using an Aurora (Cytek), and data analysis was conducted with OMIQ software (Dotmatics). For the gating strategies for flow cytometry experiments, refer to Supplementary Fig. 1.

### Cell sorting

Bulk RNA was isolated from 10^6^ cells, sorted with a MACS mouse untouched CD8^+^ T cells Isolation Kit (Miltenyi Biotec) from splenocytes, using the RNA isolation mini kit (Invitrogen) following the manufacturer’s instructions. RNA concentration was determined using NanoDrop spectrophotometer (ThermoFisher Scientific) and volumes were adjusted to ensure uniform measurement across all samples.

### RNA sequencing

For the library preparation the quality of the isolated RNA was determined with a Qubit (1.0) Fluorometer (Life Technologies) and a Fragment Analyzer (Agilent). Only those samples with a 260 nm/280 nm ratio between 1.8–2.1 and a 28S/18S ratio within 1.5–2 were further processed. The Illumina stranded mRNA Prep Ligation (Illumina) was used in the succeeding steps. Briefly, total RNA samples (100-1000 ng) were polyA enriched and then reverse-transcribed into double-stranded cDNA. The cDNA samples were fragmented, end-repaired and adenylated before ligation of pre-index anchors to the ends of double-stranded cDNA fragments. A subsequent PCR amplification step was performed to selectively amplify the anchor-ligated DNA fragments and add the index adapter sequences, resulting in dual-indexed library. The quality and quantity of the libraries were validated using Qubit (1.0) Fluorometer and the Fragment Analyzer (Agilent, Santa Clara, California, USA). The product is a smear with an average fragment size of approximately 300 bp. The libraries were normalized to 10nM in Tris-Cl 10 mM, pH8.5 with 0.1% Tween 20. After library quantification, libraries were prepared for loading accordingly to the NovaSeq workflow with the NovaSeq6000 Reagent Kit (Illumina). Cluster generation and sequencing were performed on a NovaSeq6000 System with a run configuration of single end 100bp.

### RNA analysis

For the RNAseq data analysis, first, the raw reads underwent preprocessing to ensure data quality which involved the removal of adapter sequences, trimming of low-quality ends, and filtering out reads with a Phred quality score below 20, achieved using Fastp (Version 0.23.4)^56^. Following preprocessing, the high-quality reads were pseudo-aligned to the Mouse reference transcriptome (build GRCm39) based on Gencode release M31. Quantification of gene-level expression was performed using Kallisto (Version 0.46.1)^57^. Differential expression analysis was conducted utilizing the generalized linear model implemented in the Bioconductor package DESeq2 (R version: 4.3.0, DESeq2 version: 1.40.1)^58^. Genes exhibiting altered expression levels with an adjusted p-value (Benjamini and Hochberg method) below 0.05 were considered differentially expressed.

### UMAP and PhenoGraph

Cell data first underwent preliminary analysis in OMIQ for computational analysis in flow cytometry. Clusters were then assigned using PhenoGraph^45^. The CD8+ T cell gated events were downsampled to ensure an equivalent number of cells per condition. This was followed by dimensionality reduction using Uniform Manifold Approximation and Projection (UMAP) for visualization purposes ^59^. Cluster assignments were annotated based on existing knowledge of phenotypes for CD8^+^ T cell subtypes. This approach allowed for accurate identification and comparison of various CD8^+^ T cell populations across different experimental conditions.

### Antibodies used

CD11b BUV395 (M1/70; 1:100), BD; BCL-6 BUV496 (K112-91; 1:50), BD; CD48 BUV563 (HM48-1; 1:100), BD; CXCR3/CD183 BUV 661 (CXCR3; 1:50), BD; CD44 BUV737 (IM7; 1:50), BD; KLRG1 BUV805 (2F1; 1:50), BD; TCF-1 BV421 (S33-966; 1:50), BD; PD-1 eFluor450 (J43; 1:100), eBioscience; CD25 BV480 (PC61; 1:50), BD; Ly-6C BV570 (HK1.4; 1:100), BioLegend; NK1.1 BV605 (PK136; 1:50), BioLegend; FAS/CD95 BV650 (Jo2; 1:50), BD; TIM-3 BV711 (RMT3-23; 1:50), BioLegend; CD122 BV750 (5H4; 1:50), BD; CD122 PerCP-Cy5.5 (TM-β1; 1:50), BD; MHCII BV785 (M5/114.15.2; 1:100), BioLegend; CD62L BB515 (MEL-14; 1:50), BD; Ki-67 AF532 (SolA15; 1:50), eBioscience; CD127 PerCP-Cy5.5 (A7R34; 1:50), eBioscience; NKG2D/CD314 PerCP-eFluor710 (CX5; 1:50), eBioscience; FASL/CD178 PE (MFL3; 1:50), BioLegend; CD4 PE-TexasRed (RM4-5; 1:50), Thermo Fisher Scientific; INF-γPE-AF610 (XMG1.2; 1:50), Thermo Fisher Scientific; CD11c PE-Cy5 (N418; 1:50), Thermo Fisher Scientific; GZMB PE-Cy5.5 (NGZB; 1:100), Thermo Fisher Scientific; TRAIL/CD253 PE-Cy7 (N2B2; 1:50), BioLegend; CD27 APC (LG.7F9; 1:100), Thermo Fisher Scientific; CD27 AF700 (LG.7f9; 1:50), Thermo Fisher Scientific; 2B4/CD244 AF647 (2B4.2; 1:50), BioLegend; CD3 AF700 (17A2, 1:33), Biolegend; CD8 APC-Cy7 (53-6.7; 1:50), BioLegend; T-BET BV785 (4B10; 1:50), BioLegend; TNF APC (MP6-XT22; 1:50); BioLegend.

### Statistics and graphics

OMIQ (www.omiq.ai) was used for analysis of cytometric data. Statistical analysis was performed using GraphPad (v.9 Prism Software). Statistical tests used are indicated in the figure legends. *P* < 0.05 was considered significant, whereby \*\*\*\**P* < 0.0001, \*\*\**P* < 0.001, \*\**P* < 0.01 and \**P* < 0.05, unless indicated otherwise. Unless otherwise specified, mean ± s.e.m. or quartiles, range, and mean are shown alongside scatter plots with truncated violin plots. The lower and upper hinges correspond to the first and third quartiles, respectively. Unless otherwise stated, two-tailed statistical T tests were used. Original flow cytometry data plots are depicted using OMIQ contour plot with 0.2 outlier percentile setting or scatterplot. The schematics were created with BioRender (www.BioRender.com).

## Data and materials availability

All data used in the analyses are available in the main text and supplementary materials. The RNAseq data have been deposited in the European Nucleotide Archive (ENA) under accession code PRJEB76285 (https://www.ebi.ac.uk/ena/browser/view/PRJEB76285).

## Acknowledgments

We thank the members of the Pachlopnik lab and C. Münz, P. Aichele, U. Karakus for their helpful discussions. We also extend our gratitude to the staff at the Laboratory Animal Services Center at the University Animal Hospital, and the Functional Genomic Center Zurich at the University of Zurich, for their assistance with experiments. We also thank Dr. David Fraser for copy-editing assistance. This work was supported by the Clinical Research Priority Program CYTIMM-Z of the University of Zurich (to T.M., O.B. and J.P.S.), a grant from the Research Center for Children of the University Children’s Hospital Zurich (to T.M.) and the Swiss National Science Foundation (Project number: 320030_205097) (to J.P.S.).

## Author contributions

Conceptualization: TM, OB, JPS; Methodology: TM; Investigation: TM, SM, DT, TL, SW; Visualization: TM; Funding acquisition: TM, JPS; Project administration: JPS; Supervision: SV, OB, JPS; Writing – original draft: TM, JPS.

## Extended data

**Extended Data Fig. 1.**
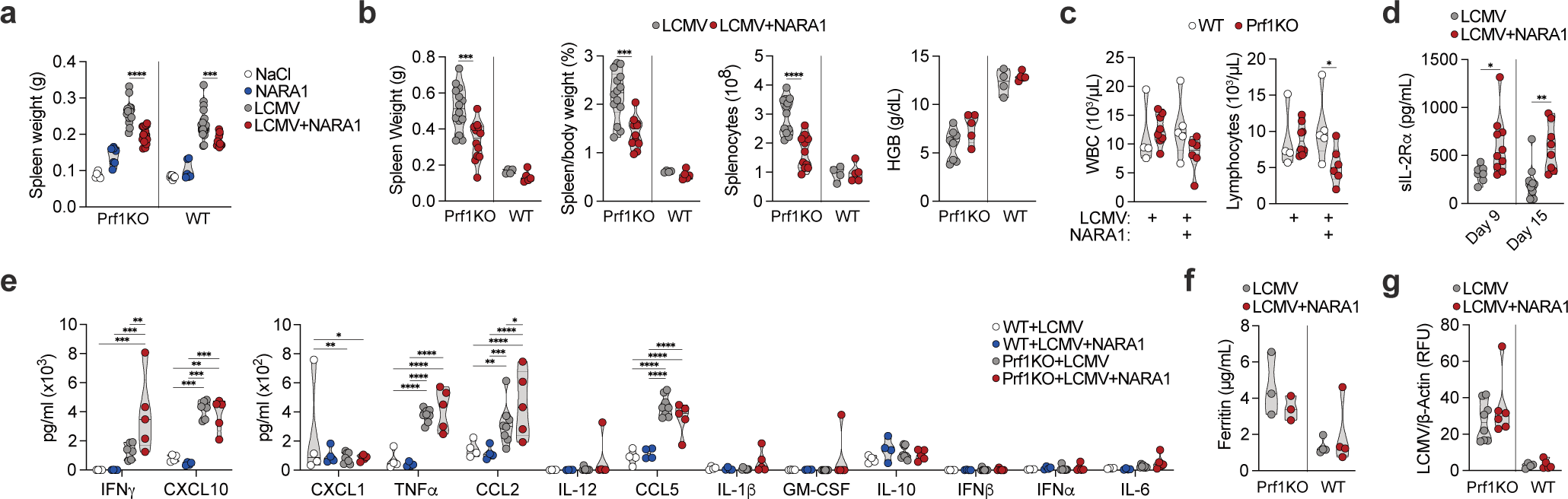
*In vivo* delivery of a CD122-biased IL-2 complex counteracts hyperinflammation and improves health in mice with an impairment in cytotoxicity. **a**, Spleen weight in mice on post-infection day 9. **b**, The variables including spleen weight, the spleen-to-mouse weight ratio, the splenocyte count, and the haemoglobin (HGB) level in mice on post-infection day 15. **c,** White blood cell (WBC) and lymphocyte counts in mice on post-infection day 15. **d**, Levels of IL-2Rαin Prf1KO mice at different times post-infection. **e**, Serum levels of cytokines involved in the mouse’s antiviral response on post-infection day 15. A two-way analysis of variance with Tukey’s correction was applied. **f**, The serum ferritin concentration on post-infection day 15. **g**, The splenic viral load on post-infection day 15. RFU=relative fluorescent unit. Each dot represents a biological replicate pooled from two or more independent experiments. n=5 to 18 per group (**a**), 4 to 14 per group (**b**), 4 to 8 per group (**c**, **e** and **g**), 7 to 10 per group (**d**), and 3 to 4 per group for (**f**). Truncated violin plots with the quartiles, range, and mean are shown alongside scatter plots. Two-tailed t-tests were applied; the statistical significance of the result is indicated by **** p<0.0001, *** p<0.001, ** p<0.01, and * p<0.05.

**Extended Data Fig. 2.**
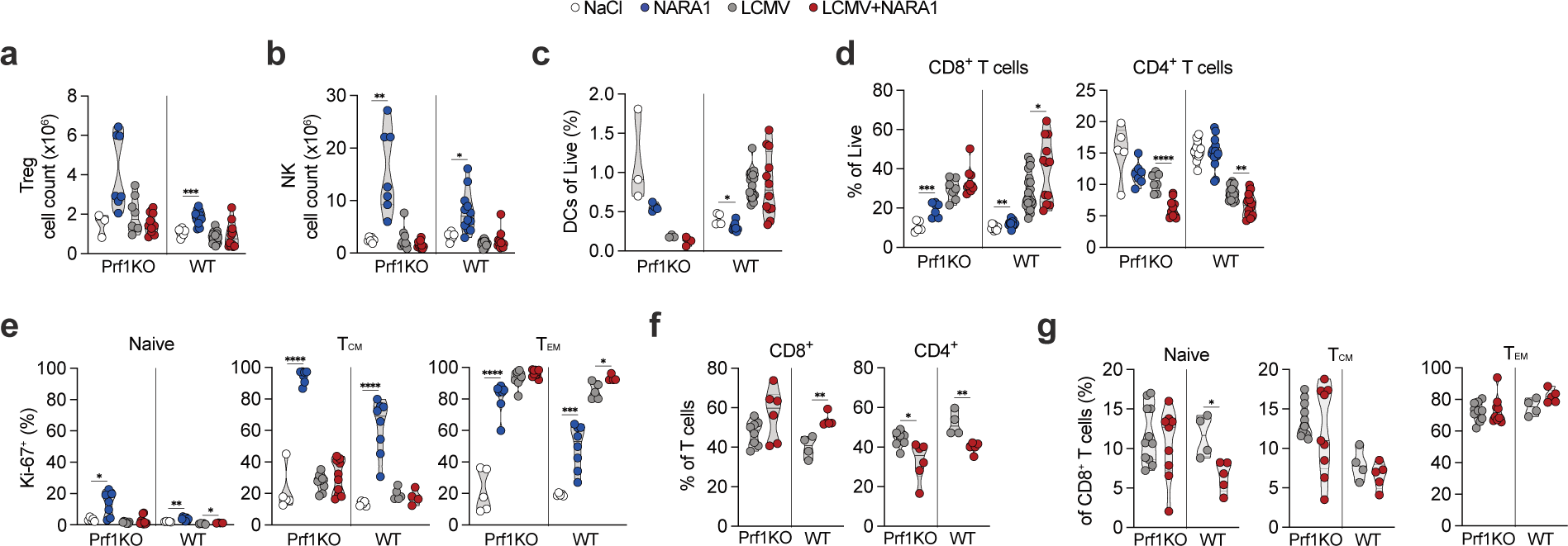
CD122-biased IL-2 delivery selectively modulates CD8 T cell differentiation during infection, independently of Prf. **a**, Total Treg counts. **b**, Total NK cell counts. **c**, DC counts as a proportion of total live cells. **d**, CD8^+^ and CD4^+^ T cell counts as a proportion of total live cells. **e**, The proportion of CD8^+^ T cells subsets expressing Ki-67: naive (CD62L^+^, CD44^−^) cells, central memory (T_CM_: CD62L^+^, CD44^+^) cells, and effector memory (T_EM_: CD62L^−^, CD44^+^) cells. **f**, Relative CD8^+^ and CD4^+^ T cell counts. **g**, Relative CD8^+^ T cell subset counts. The cells described in **a**−**d** were taken from spleens on post-infection day 9, and the cells described in the other panels were taken on post-infection day 15. Each dot represents a biological replicate pooled from two or more independent experiments. n=3 to 12 per group (**a**), 5 to 12 per group (**b**), 3 to 18 per group (**c**), 5 to 18 per group (**d**), 5 to 10 per group (**e**), 4 to 8 per group (**f**), and 4 to 11 per group (**g**). Truncated violin plots with the quartiles, range, and mean are shown alongside scatter plots. Two-tailed t-tests were applied; the statistical significance of the result is indicated by **** p<0.0001, *** p<0.001, *** p<0.01, * p<0.05 (p>0.05 in the absence of asterisks).

**Extended Data Fig. 3.**
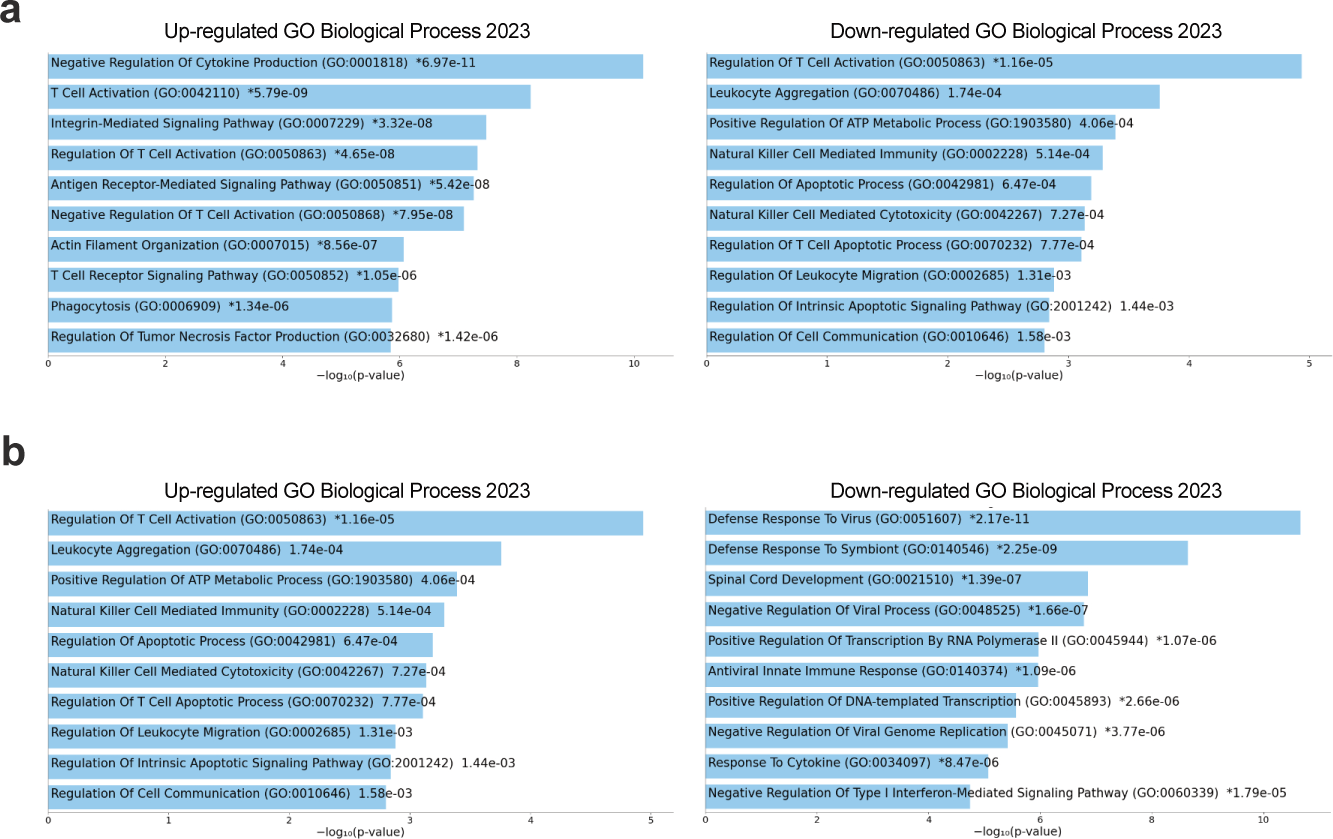
Transcriptomic analysis reveals induction of an exhausted signature in Prf-deficient CD8 T cells following IL-2 stimulation. **a**, The top 10 upregulated (left) and downregulated (right) DEGs in a Gene Ontology enrichment analysis of LCMV-infected Prf1KO mice on post-infection day 9 (after NARA1 treatment). **b**, The top 10 upregulated (left) and downregulated (right) DEGs in a Gene Ontology enrichment analysis of LCMV-infected WT mice on post-infection day 9 (after NARA1 treatment).

**Extended Data Fig. 4.**
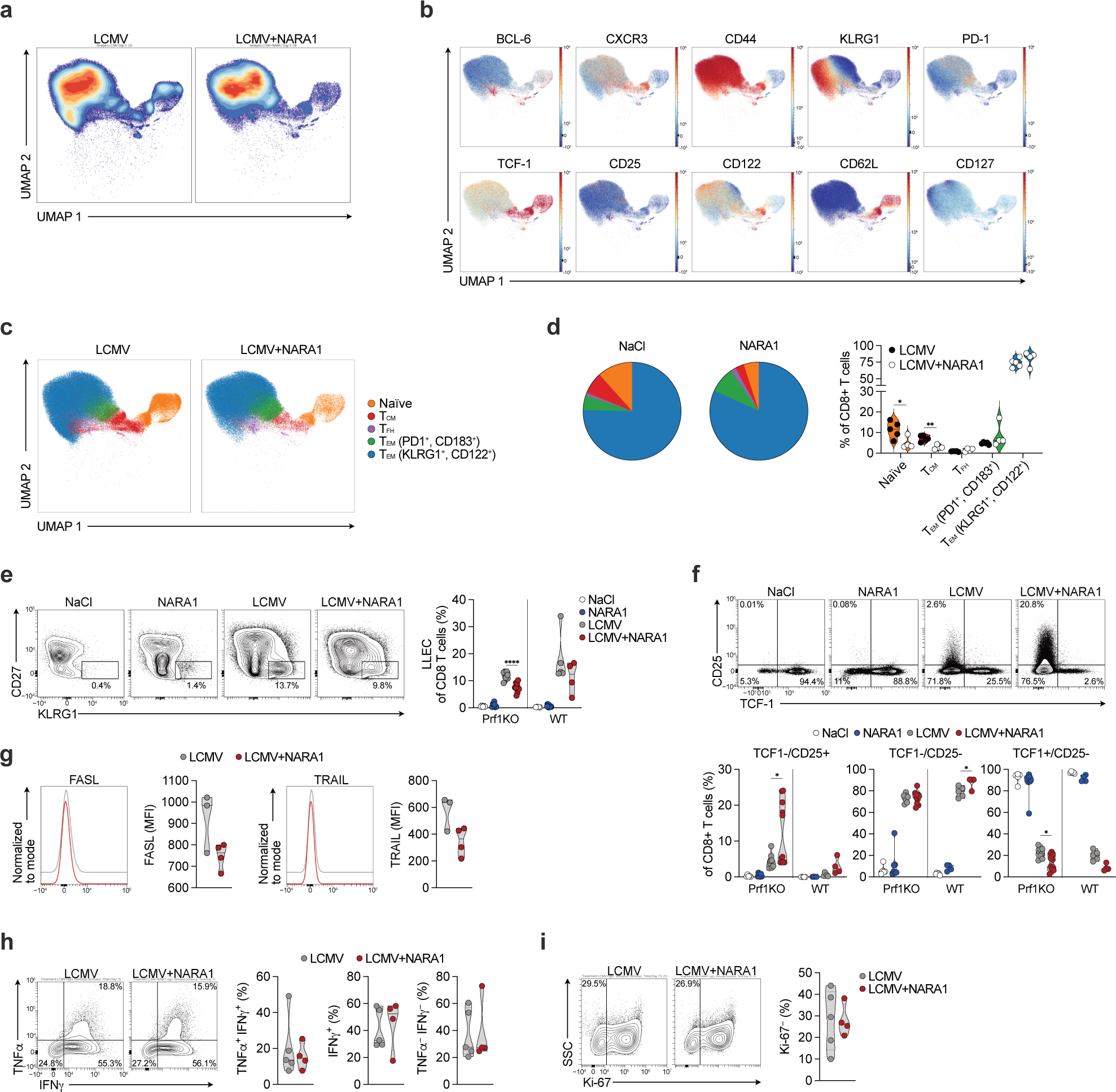
*In vivo* IL-2 treatment prompts Prf1KO CD8 T cells to adopt an exhausted phenotype and a functional state that is essential for mitigating hyperinflammation. **a**, Stacked uniform manifold approximation and projection (UMAP) plots of high-dimensional spectral flow cytometry data from gated CD8^+^ T cells obtained from the spleen of LCMV-infected WT mice after treatment with saline solution (n=5) or NARA1 (n=4) on post-infection day 9, coloured by distribution density. **b**, UMAP plots of selected markers in cells from WT mice infected with LCMV and treated with NARA1, scaled by fluorescent intensity. **c**, UMAP plots of CD8^+^ T cells from WT mice infected with LCMV, subclustered using PhenoGraph and 26 marker expression profiles. T_CM_=central memory T cells, T_PEX_=pre-exhausted T cells, T_FH_=T follicular helper cells, T_EM_=effector memory T cells. **d**, Proportions of cells, by treatment. **e**, Flow cytometry plots (left) and counts (right) of LLEC CD8^+^ T cells. **f**, Flow cytometry plots (above) and counts (below) of CD25 and TCF-1 expression on CD8^+^ T cells. **g**, FASL and TRAIL surface expression in CD8^+^ T cells from Prf1KO mice, on post-infection day 9. **h**, Flow cytometry plots and counts depicting IFNg and TNFa production by CD8^+^ T cells from spleens collected on post-infection day 15, following stimulation for 5 h. The plots were gated on CD8^+^ T cells. **i**, Flow cytometry plots and counts depicting Ki-67 expression by CD8^+^ T cells isolated from spleens, following stimulation for 5 h. The plots were gated on CD8^+^ T cells. Each dot represents a biological replicate pooled from two or more independent experiments. n= 5 to 12 per group (**e**), 5 to 10 per group (**f**), 3 to 4 per group (**g**), and 4 to 5 per group for (**h** and **i**). Truncated violin plots with the quartiles, range, and mean are shown alongside scatter plots. Two-tailed t-tests were applied; the statistical significance of the result is indicated by **** p<0.0001, *** p<0.001, ** p<0.01, and * p<0.05.

**Extended Data Fig. 5.**
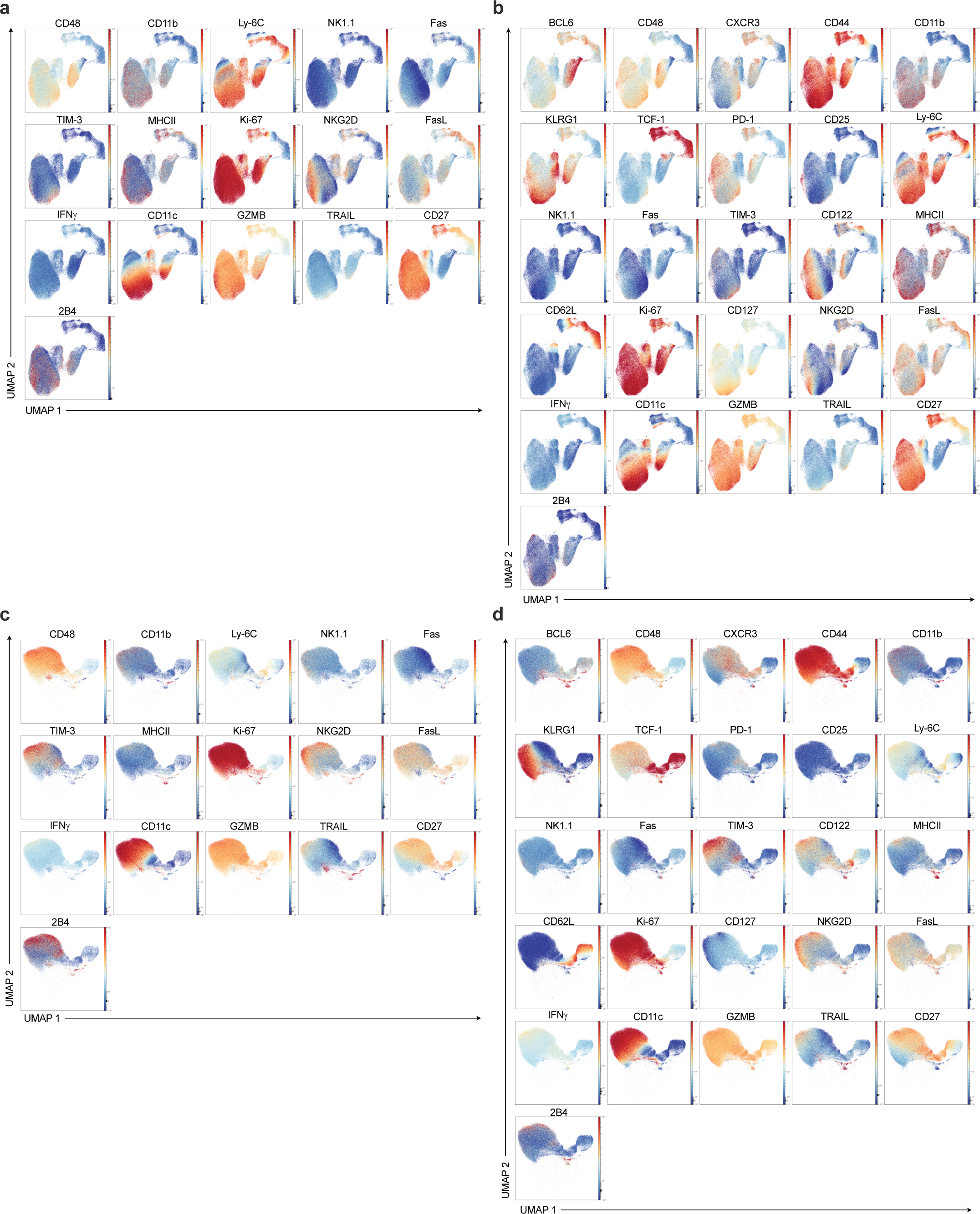
*In vivo* IL-2 treatment prompts Prf1KO CD8 T cells to adopt an exhausted phenotype and a functional state that is essential for mitigating hyperinflammation. **a**, Stacked uniform manifold approximation and projection (UMAP) plots of plots of high-dimensional spectral flow cytometry data from gated CD8^+^ T cells obtained from the spleen of Prf1KO mice infected with LCMV and treated with NARA1, scaled by fluorescent intensity (n=4). **b**, Stacked UMAP plots of all the selected markers expressed by CD8^+^ T cells from Prf1KO mice infected with LCMV (n=3). **c**, Stacked UMAP plots of selected markers in CD8^+^ T cells from WT mice infected with LCMV and treated with NARA1, scaled by fluorescent intensity (n=4). **d**, Stacked UMAP plots of all the selected markers expressed by CD8^+^ T cells from WT mice infected with LCMV (n=5).

**Supplementary Fig. 1.**
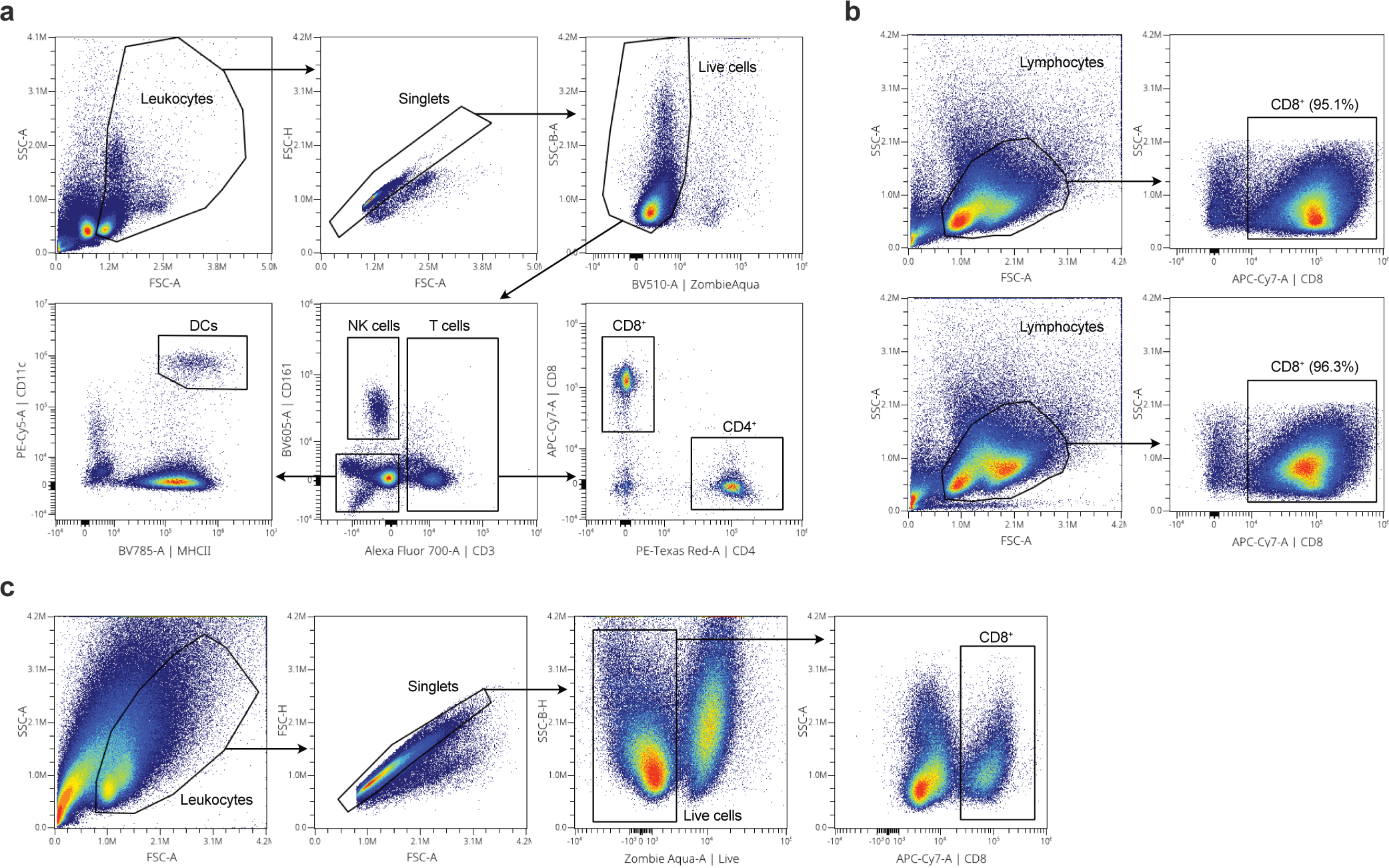
Gating strategies for flow cytometry. Gating strategies used for flow cytometry analysis of Tregs, NK cells, DCs, and CD4^+^ and CD8^+^ T cells in cell-counting experiments (**a**), cell sorting for RNAseq (**b**), or restimulation experiments (**c**).

## References

1. George, M. R. Hemophagocytic lymphohistiocytosis: review of etiologies and management. J Blood Med 5, 69–86 (2014).

2. Fajgenbaum David C. & June Carl H. Cytokine Storm. New England Journal of Medicine 383, 2255–2273 (2020).

3. Schmid, J. P. et al. Inherited defects in lymphocyte cytotoxic activity. Immunological Reviews 235, 10–23 (2010).

4. Kägi, D. et al. Cytotoxicity mediated by T cells and natural killer cells is greatly impaired in perforin-deficient mice. Nature 369, 31–37 (1994).

5. Sullivan, K. E., Delaat, C. A., Douglas, S. D. & Filipovich, A. H. Defective Natural Killer Cell Function in Patients with Hemophagocytic Lymphohistiocytosis and in First Degree Relatives. Pediatric Research 44, 465–468 (1998).

6. Stepp, S. E. Perforin Gene Defects in Familial Hemophagocytic Lymphohistiocytosis. Science 286, 1957–1959 (1999).

7. Trapani, J. A. & Smyth, M. J. Functional significance of the perforin/granzyme cell death pathway. Nat Rev Immunol 2, 735–747 (2002).

8. Badovinac, V. P., Hamilton, S. E. & Harty, J. T. Viral Infection Results in Massive CD8+ T Cell Expansion and Mortality in Vaccinated Perforin-Deficient Mice. Immunity 18, 463–474 (2003).

9. Jordan, M. B., Hildeman, D., Kappler, J. & Marrack, P. An animal model of hemophagocytic lymphohistiocytosis (HLH): CD8+ T cells and interferon gamma are essential for the disorder. Blood 104, 735–743 (2004).

10. Pachlopnik Schmid, J., et al. Neutralization of IFNgamma defeats haemophagocytosis in LCMV-infected perforin- and Rab27a-deficient mice. EMBO Mol Med 1, 112–124 (2009).

11. Jenkins, M. R. et al. Failed CTL/NK cell killing and cytokine hypersecretion are directly linked through prolonged synapse time. J Exp Med 212, 307–317 (2015).

12. Spolski, R., Li, P. & Leonard, W. J. Biology and regulation of IL-2: from molecular mechanisms to human therapy. Nat Rev Immunol 18, 648–659 (2018).

13. Rosenberg, S. A. IL-2: The First Effective Immunotherapy for Human Cancer. J Immunol 192, 5451–5458 (2014).

14. Boyman, O. & Sprent, J. The role of interleukin-2 during homeostasis and activation of the immune system. Nature Reviews Immunology 12, 180–190 (2012).

15. Chinen, T. et al. An essential role for the IL-2 receptor in Treg cell function. Nat Immunol 17, 1322–1333 (2016).

16. Hernandez, R., Põder, J., LaPorte, K. M. & Malek, T. R. Engineering IL-2 for immunotherapy of autoimmunity and cancer. Nat Rev Immunol 22, 614–628 (2022).

17. Kawakami, R. & Sakaguchi, S. Regulatory T Cells for Control of Autoimmunity. Adv Exp Med Biol 1444, 67–82 (2024).

18. Boyman, O., Kovar, M., Rubinstein, M. P., Surh, C. D. & Sprent, J. Selective Stimulation of T Cell Subsets with Antibody-Cytokine Immune Complexes. Science 311, 1924–1927 (2006).

19. Arenas-Ramirez, N., Woytschak, J. & Boyman, O. Interleukin-2: Biology, Design and Application. Trends Immunol 36, 763–777 (2015).

20. Raeber, M. E., Sahin, D. & Boyman, O. Interleukin-2–based therapies in cancer. Science Translational Medicine 14, eabo5409 (2022).

21. Wilson, M. S. et al. Suppression of Murine Allergic Airway Disease by IL-2:Anti-IL-2 Monoclonal Antibody-Induced Regulatory T Cells1. The Journal of Immunology 181, 6942– 6954 (2008).

22. Webster, K. E. et al. In vivo expansion of T reg cells with IL-2–mAb complexes: induction of resistance to EAE and long-term acceptance of islet allografts without immunosuppression. Journal of Experimental Medicine 206, 751–760 (2009).

23. Krieg, C., Létourneau, S., Pantaleo, G. & Boyman, O. Improved IL-2 immunotherapy by selective stimulation of IL-2 receptors on lymphocytes and endothelial cells. Proc Natl Acad Sci U S A 107, 11906–11911 (2010).

24. Arenas-Ramirez, N. et al. Improved cancer immunotherapy by a CD25-mimobody conferring selectivity to human interleukin-2. Science Translational Medicine 8, 367ra166-367ra166 (2016).

25. Humblet-Baron, S. et al. IL-2 consumption by highly activated CD8 T cells induces regulatory T-cell dysfunction in patients with hemophagocytic lymphohistiocytosis. J. Allergy Clin. Immunol. 138, 200–209.e8 (2016).

26. Létourneau, S. et al. IL-2/anti-IL-2 antibody complexes show strong biological activity by avoiding interaction with IL-2 receptor αsubunit CD25. Proc Natl Acad Sci U S A 107, 2171– 2176 (2010).

27. Moore, K. W., Malefyt, R. de W., Coffman, R. L. & O’Garra, A. Interleukin-10 and the Interleukin-10 Receptor. Annual Review of Immunology 19, 683–765 (2001).

28. Saraiva, M., Vieira, P. & O’Garra, A. Biology and therapeutic potential of interleukin-10. Journal of Experimental Medicine 217, e20190418 (2019).

29. Raeber, M. E., Rosalia, R. A., Schmid, D., Karakus, U. & Boyman, O. Interleukin-2 signals converge in a lymphoid–dendritic cell pathway that promotes anticancer immunity. Science Translational Medicine 12, (2020).

30. Waggoner, S. N., Cornberg, M., Selin, L. K. & Welsh, R. M. Natural killer cells act as rheostats modulating antiviral T cells. Nature 481, 394–398 (2012).

31. Sepulveda, F. E. et al. A novel immunoregulatory role for NK-cell cytotoxicity in protection from HLH-like immunopathology in mice. Blood 125, 1427–1434 (2015).

32. Pallmer, K. et al. NK cells negatively regulate CD8 T cells via natural cytotoxicity receptor (NCR) 1 during LCMV infection. PLOS Pathogens 15, e1007725 (2019).

33. Allan, R. S. et al. Migratory dendritic cells transfer antigen to a lymph node-resident dendritic cell population for efficient CTL priming. Immunity 25, 153–162 (2006).

34. Pipkin, M. E. et al. Interleukin-2 and Inflammation Induce Distinct Transcriptional Programs that Promote the Differentiation of Effector Cytolytic T Cells. Immunity 32, 79–90 (2010).

35. West, E. E. et al. PD-L1 blockade synergizes with IL-2 therapy in reinvigorating exhausted T cells. J Clin Invest 123, 2604–2615 (2013).

36. Blattman, J. N. et al. Therapeutic use of IL-2 to enhance antiviral T-cell responses in vivo. Nat Med 9, 540–547 (2003).

37. Liu, Y. et al. IL-2 regulates tumor-reactive CD8+ T cell exhaustion by activating the aryl hydrocarbon receptor. Nat Immunol 22, 358–369 (2021).

38. Badovinac, V. P., Porter, B. B. & Harty, J. T. Programmed contraction of CD8+ T cells after infection. Nat Immunol 3, 619–626 (2002).

39. Kögl, T. et al. Hemophagocytic lymphohistiocytosis in syntaxin-11–deficient mice: T-cell exhaustion limits fatal disease. Blood 121, 604–613 (2013).

40. Wherry, E. J. T cell exhaustion. Nat Immunol 12, 492–499 (2011).

41. Wherry, E. J. & Kurachi, M. Molecular and cellular insights into T cell exhaustion. Nat Rev Immunol 15, 486–499 (2015).

42. McLane, L. M., Abdel-Hakeem, M. S. & Wherry, E. J. CD8 T Cell Exhaustion During Chronic Viral Infection and Cancer. Annual Review of Immunology 37, 457–495 (2019).

43. Cornberg, M. et al. Clonal Exhaustion as a Mechanism to Protect Against Severe Immunopathology and Death from an Overwhelming CD8 T Cell Response. Front. Immunol. 4, (2013).

44. Blank, C. U. et al. Defining ‘T cell exhaustion’. Nat Rev Immunol 19, 665–674 (2019).

45. Levine, J. H. et al. Data-Driven Phenotypic Dissection of AML Reveals Progenitor-like Cells that Correlate with Prognosis. Cell 162, 184–197 (2015).

46. Tusher, V. G., Tibshirani, R. & Chu, G. Significance analysis of microarrays applied to the ionizing radiation response. Proceedings of the National Academy of Sciences 98, 5116–5121 (2001).

47. Jin, H.-T. et al. Cooperation of Tim-3 and PD-1 in CD8 T-cell exhaustion during chronic viral infection. Proceedings of the National Academy of Sciences 107, 14733–14738 (2010).

48. Escobar, G., Mangani, D. & Anderson, A. C. T cell factor 1 (Tcf1): a master regulator of the T cell response in disease. Sci Immunol 5, eabb9726 (2020).

49. Renkema, K. R. et al. KLRG1+ Memory CD8 T cells Combine Properties of Short-lived Effectors and Long-lived Memory. J Immunol 205, 1059–1069 (2020).

50. Koreth John et al. Interleukin-2 and Regulatory T Cells in Graft-versus-Host Disease. New England Journal of Medicine 365, 2055–2066 (2011).

51. Sun, J., Madan, R., Karp, C. L. & Braciale, T. J. Effector T cells control lung inflammation during acute influenza virus infection by producing IL-10. Nat Med 15, 277–284 (2009).

52. Sahin, D. et al. An IL-2-grafted antibody immunotherapy with potent efficacy against metastatic cancer. Nature Communications 11, 6440 (2020).

53. Beltra, J.-C. et al. IL2Rβ-dependent signals drive terminal exhaustion and suppress memory development during chronic viral infection. Proceedings of the National Academy of Sciences 113, E5444–E5453 (2016).

54. Kalia, V. et al. Prolonged Interleukin-2RαExpression on Virus-Specific CD8+ T Cells Favors Terminal-Effector Differentiation In Vivo. Immunity 32, 91–103 (2010).

55. Hashimoto, M. et al. PD-1 combination therapy with IL-2 modifies CD8+ T cell exhaustion program. Nature 610, 173–181 (2022).

56. Chen, S., Zhou, Y., Chen, Y. & Gu, J. fastp: an ultra-fast all-in-one FASTQ preprocessor. Bioinformatics 34, i884–i890 (2018).

57. Bray, N. L., Pimentel, H., Melsted, P. & Pachter, L. Near-optimal probabilistic RNA-seq quantification. Nat Biotechnol 34, 525–527 (2016).

58. Love, M. I., Huber, W. & Anders, S. Moderated estimation of fold change and dispersion for RNA-seq data with DESeq2. Genome Biology 15, 550 (2014).

59. Liechti, T. et al. An updated guide for the perplexed: cytometry in the high-dimensional era. Nat Immunol 22, 1190–1197 (2021).

